# Knockdown of the long non-coding RNA DNAAF3-AS1 activates the p53 pathway and remodels the H3K36me3 landscape

**DOI:** 10.64898/2025.12.15.694275

**Authors:** Anna Budkina, Anatoliy Zubritskiy, Daria Marakulina, Andrey Buyan, Ivan Kulakovsky, Chi Wai Yip, Jude Aneke, Jay Shin, Yulia A. Medvedeva

## Abstract

Long non-coding RNAs are increasingly recognised as chromatin regulators, yet most remain functionally uncharacterised. *DNAAF3-AS1*, transcribed antisense to the ciliary dynein assembly factor gene *DNAAF3*, is almost exclusively associated with H3K36me3 in the HiMoRNA database. H3K36me3 ChIP-seq in human dermal fibroblasts following *DNAAF3-AS1* knockdown with two independent antisense oligonucleotides revealed a marked reduction of gene-body H3K36me3, with residual signal shifted towards promoters. In FANTOM6 CAGE, *DNAAF3-AS1* knockdown activated the p53 pathway and triggered differential promoter usage in genes enriched for p53 and apoptotic regulators. Consistently, genes that gained gene-body H3K36me3 upon knockdown were enriched for apoptosis-related pathways. In line with high expression of *DNAAF3-AS1* in testis, analysis based on the adult human testis single-cell atlas suggests that *DNAAF3-AS1* may contribute to promoter-level H3K36me3 regulation during late spermatogenesis. Together, these results identify *DNAAF3-AS1* as a chromatin-associated lncRNA that links H3K36me3 maintenance to p53 signalling.

## Introduction

Human cells transcribe long non-coding RNAs (lncRNAs) in numbers that rival those of protein-coding genes [9], yet their functional annotation remains notoriously difficult. Low expression, tissue-restricted profiles, and patchy evolutionary conservation have long confounded efforts to assign roles to these transcripts [8, 26].However, what endures is often structural or positional: synteny with neighboring genes, short sequence motifs, and secondary structure are frequently preserved [18], hinting at latent function [26]. Indeed, mounting evidence positions lncRNAs as tightly regulated players in diverse molecular mechanisms [4, 41].

*DNAAF3-AS1* (*CTD-2587H24.5*, *AC010327.4*, *ENSG00000267577*) exemplifies this class. Classified as a transcribed enhancer lncRNA via ChIA-PET network analysis, it is thought to operate through chromatin-associated regulatory modules [59]. Loss-of-function studies further cast it as a cis-acting repressor, with its target genes overrepresented within its topologically associated domain [17]. Its antisense neighbor, an axonemal dynein assembly factor *DNAAF3* is required for the cytoplasmic pre-assembly of axonemal dynein arms, loss of *DNAAF3* function depletes both outer and inner dynein arms from cilia and causes primary ciliary dyskinesia [44]. In *Drosophila*, the *DNAAF3* orthologue is expressed in cells bearing motile cilia and is required for sperm motility [77]. Mechanistically, *DNAAF3* resembles S-adenosylmethionine-dependent methyltransferases and methylates dynein heavy chains at critical residues [53]. Consistently, *DNAAF3* mutations also drive severe asthenozoospermia, teratospermia, and male infertility [10, 61].

LncRNAs frequently interact with chromatin, influencing epigenetic regulation at specific loci and across broader chromosomal architecture [23, 39, 46, 48]. LncRNAs can shape histone modification patterns by guiding or scaffolding chromatin-modifying complexes or acting indirectly on chromatin-modifying enzymes expression for different histone modifications [21, 47, 67, 73] including H3K36me3 [34, 66]. H3K36me3 is deposited by the histone methyltransferase *SETD2*, which is recruited to elongating RNA polymerase II through its phosphorylated C-terminal domain [64] and through the elongation factors *SPT6* and *IWS1* [40, 68]. Consequently, H3K36me3 is propagated along gene bodies and is preferentially enriched at exons and towards the 3 ends of genes [57]. The processes of establishing H3K36me1 and H3K36me2 are mainly regulated by the *NSD1*, *NSD2*, *NSD3* and *ASH1L* proteins independently of *SETD2* activity. However, there is evidence that loss of function of *NSD* proteins may influence H3K36me3 levels [28]. In promoters, *SMYD5* deposits H3K36me3 specifically at promoter regions [71]. H3K36me3 was also identified in atypical heterochromatin simultaneously enriched with H3K9me3, while with removal of *SETDB1*, the H3K9me3 writer protein, such domains lose both marks and gain signatures of active enhancers [6]. The H3K36 methylation is enzymatically removed from chromatin by members of histone demethylase protein families containing the Jumonji domain, such as *KDM4A*, *KDM4B* and *KDM4C* [54].

We previously built HiMoRNA [42], a multi-omics resource that correlates lncRNA expression with histone modification peaks to computationally infer epigenetic effects, and later supported it with RNA-chromatin interactome data to validate lncRNA-mediated epigenetic regulation [27]. Here, integrating these frameworks and leveraging HiMoRNA’s correlations between *DNAAF3-AS1* expression and histone mark intensity, we set out to dissect its functional repertoire and epigenetic regulatory logic.

## Materials and Methods

### ASO transfection protocol

Primary human dermal fibroblasts (Lonza, catalog number: C2509) were maintained in DMEM supplemented with 10% FBS at 37°C with 5% CO. In order to knockdown *DNAAF3-AS1* (*CTD-2587H24.5*), two GapmeR antisense oligonucleotides (ASOs) designed against distinct regions (Exiqon) were selected as biological replicates based on consistent transcriptomic profiles after knockdown (Pearson’s r *>* 0.98 between replicates) and strong knockdown efficiency observed from a previous study [49]. CTD-2587H24.5-ASO_G0267577_01 5’-AGGAGGCTTCGTGGTG-3’ (ASO_01) CTD-2587H24.5-ASO_G0267577_03 5’-TAGGACAGATCTGCGG-3’ (ASO_03). DNAAF3-AS knockdown was performed as previously described [49]. Briefly, cells were transfected with a final concentration of 2 nM ASOs using Lipofectamine RNAiMAX (Thermo Fisher 13778150), in parallel with a non-targeting ASO as a negative control (NC_A 5’-AACACGTCTATACGC-3’ (Exiqon)). At 48 hours post-transfection, RNA was extracted by RNeasy (Qiagen). Knockdown efficiency was validated by qRT-PCR using specific primers: CTD-2587H24.5 Fw 5’-TATGGGGCCTCAACACATGC-3’ CTD-2587H24.5 Rev 5’-TCAGACAGAGGAGGCTTCGT-3’. Compared to the NC_A control, the two ASOs showed 77-84% knockdown of DNAAF3-AS1.

### Chromatin immunoprecipitation and sequencing

Chromatin immunoprecipitation followed by high-throughput sequencing was performed to investigate histone mark distribution in human dermal fibroblasts (HDF) with and without lncRNA knockdown. The ChIP was conducted using the “Cross-linking Chromatin Immunoprecipitation protocol” (www.abcam.com) with some modifications. 2 ug of Anti-H3K36me3 (Abcam 9050) antibodies per sample were used for chromatin immunoprecipitation. The input controls were similarly processed without any antibody. The recovered ChIP and input DNA were used to construct Illumina sequencing libraries. Sequencing was carried out with 2 biological replicates for control with non-target ASO and 2 biological replicates for knockdown (one per ASO).

### ChIP-seq data processing

Raw ChIP-Seq reads were processed with Trim Galore (v0.6.10) [30], and clean reads were mapped to chr1-22, chrX, and chrY sequences from the Homo sapiens reference genome (assembly GRCh38.p14) using Bowtie2 (v2.3.4.1) software [33] in local mode. Samtools (v1.8) [15] was used to sort bam alignment files by genomic coordinates and remove PCR duplicates. MACS3 (v3.0.0a6) [72] was used for peak calling, comparing each H3K36me3 ChIP sample against its respective input control. Peaks were identified with the following parameters: –broad and –broad-cutoff 0.1. For each ChIP sample, cross-correlation quality metrics and the fragment-length estimate were computed with phantompeakqualtools [32]. Differential binding analysis was performed using DiffBind (v3.12.0) [51]. A consensus peak set was derived by retaining every interval present in at least one sample (minOverlap = 1), and read counts were computed using dba.count with summits = FALSE. Read counts were normalised using the DESeq2 median-of-ratios method, and differential binding was tested with DESeq2 [37]; results with Benjamini–Hochberg FDR *<* 0.05 were used for the further analysis. Differential peaks were annotated using ChIPseeker (v1.38.0) [62] against the GENCODE v49 annotation, with a promoter region defined as 1,000 to +100 bp from TSS. Averaged metagene profiles were constructed using deeptools software [50] for genes with mean TPM expression above the median in human dermal fibroblasts across FANTOM6 control samples for RPKM-normalised H3K36me3 coverage. For *DNAAF3-AS1* knockdown ASO_01 and ASO_03, and two controls, genes were grouped into eight clusters by the shape of their H3K36me3 metagene profile using the k-means option of deepTools.

### Differential expression and gene set enrichment analysis

Differential gene expression results for two *DNAAF3-AS1* (*CTD-2587H24.5* in FANTOM6) knockdown experiments ASO_01 and ASO_03 were downloaded from the FANTOM6 data repository [1].Additionally, *DNAAF3-AS1*, *DNAAF3* and *SYT5*, per-library CAGE expression (TPM) from the FANTOM6 was compared between *DNAAF3-AS1* knockdown (n = 4;ASO_01 and ASO_03 pooled) and matching non-targeting controls (n = 8) that corresponded to *DNAAF3-AS1* knockdown using a two-sided Wilcoxon rank-sum test per gene. For each ASO, all tested genes were ranked by decreasing log2 fold change and subjected to gene set enrichment analysis (GSEA) of GO:BP terms with clusterProfiler (v4.13.0) [69] and the org.Hs.eg.db annotation database (v3.18.0). GO over-representation analysis was additionally performed separately for significantly upregulated and downregulated genes (FDR *<* 0.05). In both analyses, p-values were adjusted for multiple testing using the Benjamini–Hochberg procedure, and terms with adjusted p-value *<* 0.05 were considered significant. Genes with differential promoter (geneFDR *<* 0.05 and prmtrPv *<* 0.05) usage in a coding region were submitted to Metascape [3, 75] for enrichment analysis. The same procedure was applied to genes annotated with H3K36me3 peaks gained upon *DNAAF3-AS1* knockdown (FDR *<* 0.05, logFC *>* 0) in an intron or exon.

### Pathway activity inference

Pathway activities were inferred with decoupleR (v2.9.7) [5] using the PROGENy model of human signalling pathways, restricted to the 500 most responsive target genes per pathway (get_progeny(organism = “human”, top = 500)). For each ASO, each gene was assigned a signed significance statistic, sign(log2FC) × log10(FDR). This gene-level statistic vector was used as input to the multivariate linear model method (run_mlm); pathways with fewer than five measured targets were excluded (minsize = 5).

### Differential promoter usage analysis

Promoter-level expression for the *DNAAF3-AS1* ASO knockdowns was obtained from FANTOM6 CAGE data. Starting from the TMM-normalised batch-corrected CAGE promoter expression matrix, counts per million (CPM) were extracted for each ASO knockdown library and compared against the panel of 143 non-targeting negative-control ASO libraries. Values of the normalised matrix were back-transformed from log scale to the linear CPM scale and were supplied to edgeR (v3.28.1) [11], promoters expressed 2 CPM in 2 or more samples were left. Differential promoter usage was then tested with diffSpliceDGE from the edgeR package: each CAGE promoter was treated as an “exon” nested within its host gene, followed by gene-level F-statistic estimation and p-value adjustment with Benjamini–Hochberg procedure. For each ASO, genes undergoing a significant differential promoter usage were defined as those with geneFDR *<* 0.05. Gene-body coordinates were taken from the GENCODE v49 annotation, and exact coding coordinates were retrieved from Ensembl through biomaRt (genomic_coding_start/genomic_coding_end), collapsed per gene to the full coding span (minimum coding start to maximum coding end across all coding transcripts). Genes with promoter differential usage in coding region were defined as genes with gene-level FDR *<* 0.05 that contained at least one promoter with prmtrPv *<* 0.05 located within the coding region of the gene. Coding region coordinates were obtained with the cdsBy function of the GenomicFeatures package (1.54.3) from the GENCODE v49 annotation.

### HiMoRNA data analysis

Associations between *DNAAF3-AS1* and H3K36me3 were obtained from the HiMoRNA database (last updated 24 April 2024; [2, 42]). In HiMoRNA, associations are defined by the Spearman correlation coefficient (SCC) between lncRNA expression and histone modification signal at individual peaks, which for H3K36me3 was calculated across 52 samples; only correlations significant after Benjamini–Hochberg correction for each lncRNA (FDR *<* 0.05) are reported. We downloaded all *DNAAF3-AS1* associations with |SCC| *>* 0.5 and retained the H3K36me3-associated entries. Peaks were annotated using the same procedure as for the ChIP-seq peaks.

### RNAChromDB colocalization analysis

Genome-wide contacts of *DNAAF3-AS1* (*AC010327.4* in database) with chromatin were downloaded from RNA-Chrom database [52] as a set of target genes for all iMargi, GRID and Red-C experiments available for Homo sapiens and pooled across experiments. RNA-chromatin target gene regions that are no more than 10 Kb apart were combined into a single set of genomic intervals. The spatial relationship between lost peaks obtained from differential binding analysis of ChIP-seq data (logFC *<* 0, p-value *<* 0.05) as query and RNA-chromatin interaction regions as reference was assessed with the GenometriCorr package (v 1.1.24) [20] using GenometriCorrelation, treating the genome as a single whole (awhole.only = TRUE) and evaluating significance against 500 random permutations of interval positions, with the alternative hypothesis set to attraction. The empirical cumulative distribution of relative distances was compared with the uniform expectation (y = 2x), and the significance of the departure was quantified by the permutation-based p-value of the ECDF deviation area (relative.distances.ecdf.deviation.area.p.value).

### Single-cell analysis

Raw single-cell counts for adult human testis were obtained from GEO accession GSE112013 and analysed with Seurat (v5.1.0) [24].The count matrix generated by the preprocessing described in Guo et al. [22]. using Cell Ranger v1.2.1 was used for downstream analysis, cells were retained if they expressed more than 500 genes and had fewer than 20% of reads mapped to the mitochondrial genome. Counts were normalised using the LogNormalize method with a scale factor of 10,000, and the 2,000 most variable genes were identified by the variance-stabilising transformation (vst) method. Cells were embedded in two dimensions by UMAP using the first 12 principal components. Clustering was carried out on a shared nearest-neighbour graph built from the first 12 principal components, followed by Louvain community detection at resolution 1, which yielded 21 clusters. Clusters were manually annotated into germ-cell and somatic populations based on the set of marker genes described in the initial paper [22]. HiMoRNA peaks associated with *DNAAF3-AS1* and H3K36me3 were partitioned by genomic context into promoter peaks and gene-body peaks (any exonic or intronic annotation) based on ChIPseeker annotation, yielding, for each context, a positive-correlation and a negative-correlation gene list. For each genomic context, the positively and negatively correlated gene lists were used as input for to AddModuleScore with default parameters, producing per-cell module scores.

## Results

### DNAAF3-AS1 knockdown activates the p53 pathway

*DNAAF3-AS1* is positioned on chromosome 19 in the human genome and is transcribed antisense to *DNAAF3* across most of its annotated isoforms, with its longest variant additionally overlapping *SYT5* (Figure 2A). This genomic architecture is evolutionarily conserved: its murine counterpart, Gm15873, is also transcribed antisense relative to Dnaaf3 on chromosome 7 (Figure S1A).Exon 2 of human *DNAAF3-AS1* overlaps a *DNAAF3* exonic region and retains 89% sequence identity with mouse Gm15873 exon 1 (BLASTN, 119/134, E-value = 11037). Across human tissues, *DNAAF3-AS1* expression is highest in testis, pituitary, fibroblasts, pancreatic islets, brain and heart (left ventricle), and *DNAAF3* is detected in a partially overlapping set of tissues (Figure 2B, Figure S1B; [14]). The relative abundance of the two transcripts nevertheless varies between tissues: in cultured fibroblasts, for example, *DNAAF3-AS1* is readily detected whereas *DNAAF3* is nearly absent, and the ratio differs again in testis. The conserved antisense arrangement, together with this partially overlapping but non-identical expression pattern, raises the question of whether the two genes are functionally coupled.

**Figure 1.**
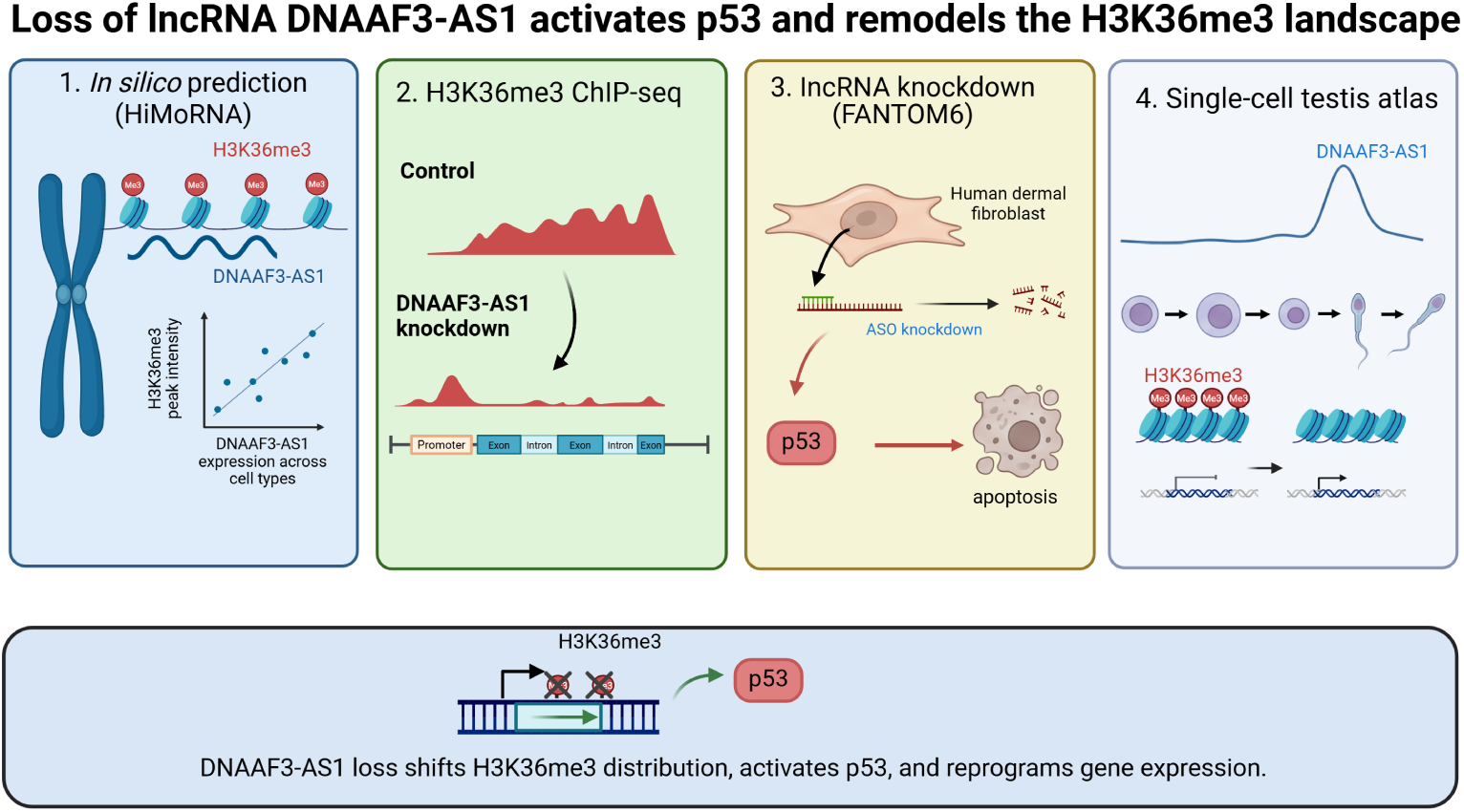
Created in BioRender. Budkina, A. (2026) https://BioRender.com/w74h360

**Figure 2.**
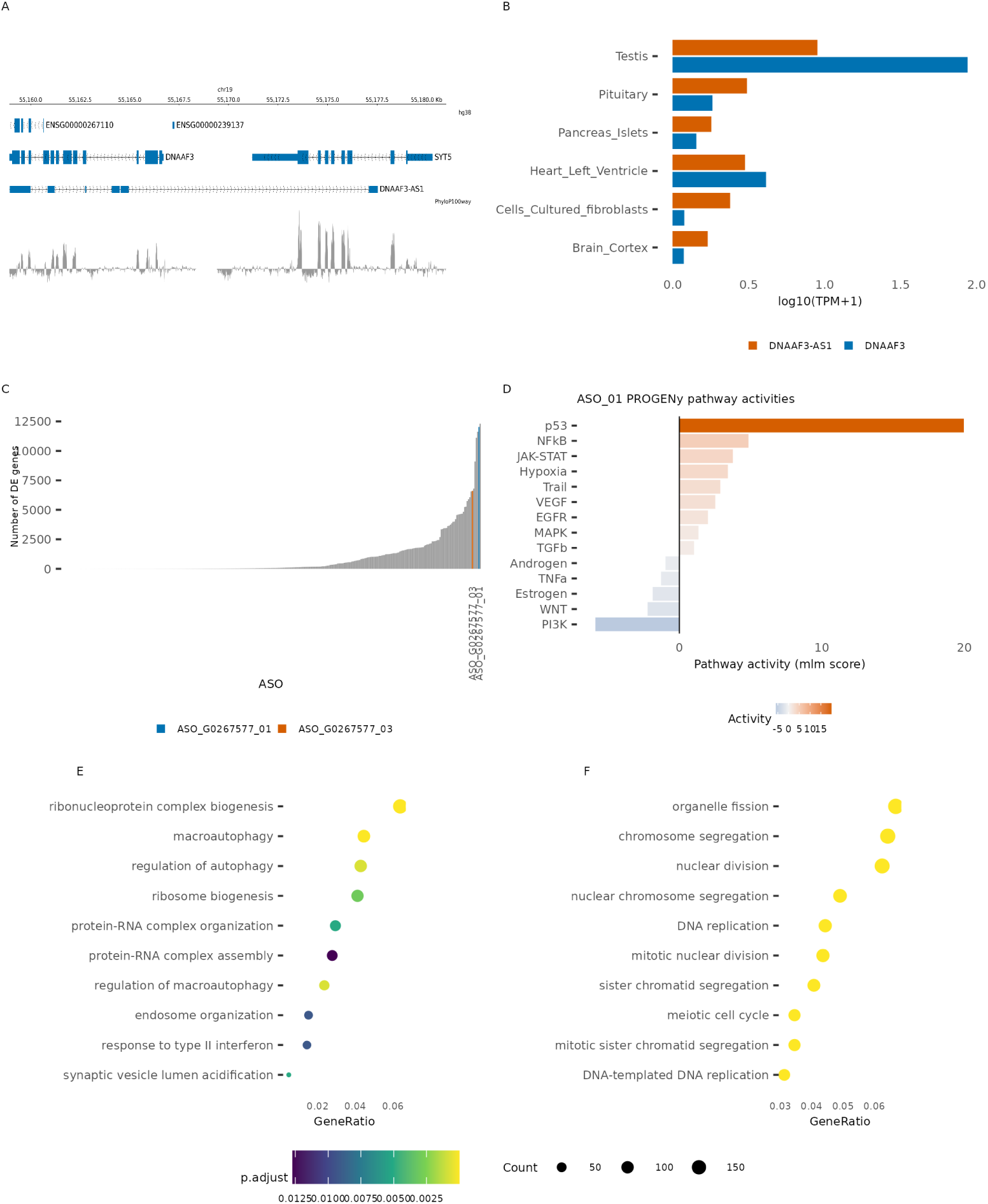
Characterisation and knockdown of the DNAAF3-AS1 locus. **A** Genomic track of the DNAAF3/DNAAF3-AS1 locus (hg38), PhyloP100-way track is shown below. **B** Median expression (log10(TPM+1)) across samples within each tissue of DNAAF3-AS1 and DNAAF3 in the top 6 GTEx v11 [14] tissues with the highest DNAAF3-AS1 expression. **C** Number of significantly differentially expressed genes (FDR *<* 0.05) per ASO in the FANTOM6 knockdown, the two DNAAF3-AS1 targeting ASOs, ASO_01 and ASO_03, are highlighted. **D** PROGENy pathway activities from the ASO_01 differential expression signature (FDR *<* 0.05). **E,F** GO biological process terms enriched among genes up-regulated (E) and down-regulated (F) upon ASO_01 knockdown

To address this question, we analysed lncRNA knockdown experiments profiled in the FANTOM6 project [49].*DNAAF3-AS1* was targeted in two independent experiments, each using a distinct antisense oligonucleotide. *DNAAF3* expression in fibroblasts was in the bottom 5% of genes. Compared with control samples transfected with non-targeting ASOs, *DNAAF3* expression increased 1.74-fold in ASO_01 knockdown and 2.99-fold in ASO_03 knockdown (Wilcoxon test p-value *<* 0.05 for ASO_01 and ASO_03 values merged, Figure S1C). In both ASOs, the number of differentially expressed genes (DEGs) ranked *DNAAF3-AS1* knockdown among the top 10 most disruptive knockdowns across all 370 ASO experiments for 169 lncRNAs tested (FDR *<* 0.05, Figure 2C, Supplementary Table 1A,B). The overlap between the two DEG sets was asymmetric: 74% of ASO_03 DEGs were also differentially expressed with ASO_01, whereas the shared genes accounted for only 36% of all DEGs (Figure S1D). This asymmetry is consistent with the knockdown efficiencies reported in FANTOM6 (95.6% for ASO_01 versus 66.8% for ASO_03) [49]: the more efficient ASO_01 produced a larger set of DEGs that encompassed most of the ASO_03 response. Pathway activity inference pointed to p53 signaling activation (Figure 2D, Figure S2A,B, Supplementary Table 1C,D),with upregulation of canonical p53 effectors *TP53INP1*, *TP53I3*, *SESN1*, *FDXR*, and *RRM2B*.

Because p53 activation is a documented sequence- and chemistry-dependent toxic effect of gapmer ASOs [55], independent of knockdown of the intended target, we benchmarked the p53 pathway activity scores of the *DNAAF3-AS1* knockdowns against those of all ASO experiments in FANTOM6. The p53 activity scores of ASO_01 and ASO_03 fell in the 98.4th and 93.5th percentiles of this distribution, respectively (Figure S2C). The p53 response to *DNAAF3-AS1* knockdown therefore exceeds that observed for the large majority of ASOs in the dataset, and its reproducibility across two independent ASOs with distinct sequences argues against a sequence-specific off-target effect, although we cannot exclude a contribution from ASO-related toxicity.

Genes upregulated in knockdown (log2FC *>* 0, FDR *<* 0.05) were enriched for autophagy regulation and positive regulation of apoptotic process (q-value *<* 0.05,) for both ASO experiments, downregulated genes (log2FC *<* 0, FDR *<* 0.05) were enriched for terms linked to a downregulation of cell-cycle genes consistent with cell cycle arrest (Figure 2E,F, Supplementary Table 1E,F). These results were supported by GSEA where cell cycle-related gene sets were negatively enriched for both ASOs (q-value *<* 0.05, Supplementary Table 1G,H).

The morphological and growth phenotypes reported in FANTOM6 also differed between the two ASOs. ASO_01 caused significant growth inhibition, an increase in cell extent and a decrease in eccentricity, changes consistent with cell rounding (Figure S3A). This combination of reduced proliferation and cell rounding is consistent with apoptosis, although it is not specific to it. ASO_03 produced no comparable change in growth rate and fewer morphological alterations. The stronger phenotype of ASO_01 thus parallels its higher knockdown efficiency and broader transcriptional response; incomplete knockdown by ASO_03 may explain its weaker effect.

### Differential promoter usage within coding regions enriches for p53 pathway and apoptotic regulators

CAGE-seq profiling in FANTOM6 permits per-gene quantification of differential promoter usage events induced by lncRNA knockdown. We selected genes with significant differential promoter usage (gene-level F-test FDR *<* 0.05). Differential promoter usage and differential expression overlapped considerably; for ASO_01, 1489 of 1960 genes with significant differential promoter usage (FDR *<* 0.05) were also differentially expressed (FDR *<* 0.05; 76.0%), compared with 692 of 1392 genes for ASO_03 (49.7%) (Figure 3A, , Supplementary Table 2A,B).

**Figure 3.**
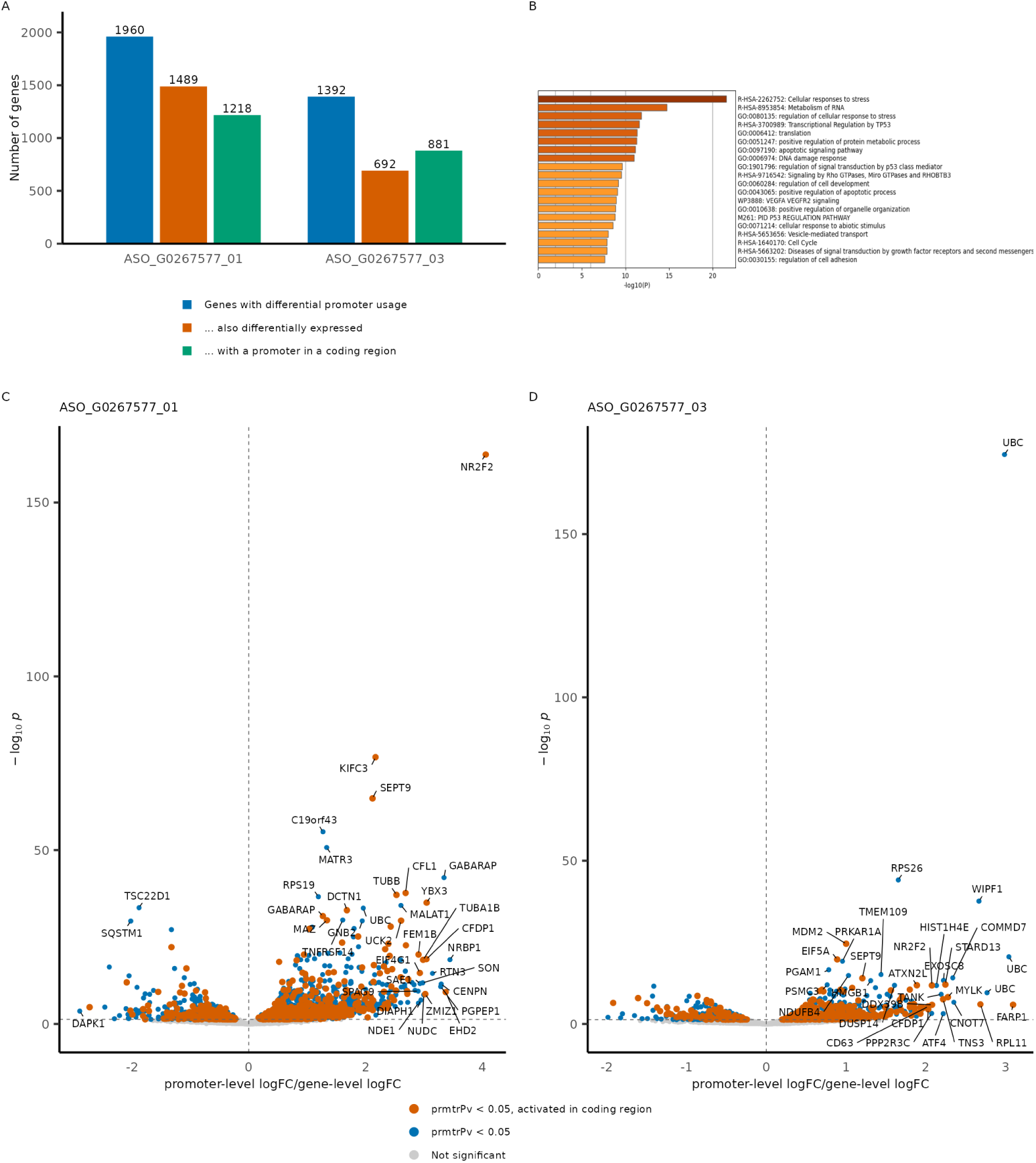
DNAAF3-AS1 knockdown triggers promoter activation in coding region. **A** Number of genes affected by differential promoter usage (gene-level FDR< 0.05) for the two DNAAF3-AS1-targeting antisense oligonucleotides (ASO_01 and ASO_03) **B** Metascape functional enrichment summary of the genes undergoing differential promoter usage upon DNAAF3-AS1 knockdown both in ASO_01 and ASO_03. **C,D** Volcano plot of promoter-level changes for ASO_01 (C) and ASO_03 (D).

We next restricted the analysis to genes in which the promoter activated or repressed upon knockdown was located within the coding region, and could therefore affect the encoded protein (Supplementary Table 2C,D). We also limited this analysis to significant effects observed in both ASOs. Functional enrichment of this subset highlighted the p53 signaling pathway and apoptosis, including cellular response to stress, transcriptional regulation by TP53, and DNA damage response(Figure 3B). Within this subset, we recovered the known differential promoter usage at *MDM2* (Figure S3B), from the constitutive P1 promoter to the p53-responsive internal P2 promoter located in intron 1, which produces a transcript with an alternative first exon [12, 70]. The recovery of this well-characterised, p53-dependent differential promoter usage serves as a positive control, indicating that the enriched signal reflects genuine activation of the p53 program.

*NR2F2*, *FASTK* and *CFDP1* were among the genes with the strongest differential promoter usage in both ASO knockdowns (Figure 3C,D, Figure S3C-E). Their connections to the p53/apoptosis programme differ in both directness and mechanism. *NR2F2*, which product acts as a context-dependent tumour promoter or suppressor in cancer [36], showed activation of intronic promoters P010 and P011, generating an isoform lacking the first exon. These promoters lie near the host gene of miR-1469 which promotes apoptosis [65]. A similar first-exon-skipping isoform, transcribed from an upstream TSS and lacking the N-terminal DNA-binding domain, has been linked to metastatic progression in melanoma [16]. *FASTK* encodes a mitochondrial RNA-binding protein activated during Fas-mediated apoptosis and involved in splicing of Fas pre-mRNA via TIA-1 [60]. *FASTK* mitochondria-specific isoform from internal translation initiation has been described previously [29]. Upon *DNAAF3-AS1* knockdown *FASTK* showed activation of a shorter first-exon-skipping isoform. *CFDP1* depletion disrupts chromatin organisation, condensin recruitment, and cell-cycle progression; only the alternative isoforms caused by differential final-exon inclusion have been previously reported for this gene, and no intronic promoter activation has been described [43].

### DNAAF3-AS1 knockdown remodels the H3K36me3 landscape

To assess whether *DNAAF3-AS1* expression is linked to the epigenetic landscape, we turned to HiMoRNA [42], where lncRNA expression is correlated with ChIP-seq peak intensity across ten histone marks (H3K27ac, H3K27me3, H3K36me3, H3K4me1, H3K4me2, H3K4me3, H3K9ac, H3K9me3, H3K79me2, H4K20me1). For *DNAAF3-AS1*, almost all correlated peaks corresponded to a single mark: 1,214 of 1,218 peaks with absolute Spearman rho *>* 0.5 mapped to H3K36me3 (Figure S4A). H3K36me3 positively and negatively correlated peaks (absolute rho *>* 0.5) showed distinct genomic distributions (Figure 4A, Supplementary Table 3). Positively correlated peaks were more frequently located in introns (first introns, 22.4% versus 16.0%; other introns, 29.5% versus 23.6%) and at promoters (26.0% versus 21.5%). Negatively correlated peaks were more frequently annotated as distal intergenic (24.0% versus 10.0%) and were enriched 5–10 kb and 10–100 kb upstream of transcription start sites (Figure S4B). These opposing distributions point to a connection between *DNAAF3-AS1* expression and H3K36me3 deposition, though whether this connection is causal, and in which direction, remains unresolved. *DNAAF3-AS1* may actively guide the H3K36me3 writing machinery to gene bodies; it may promote transcription, with H3K36me3 deposited co-transcriptionally; both the lncRNA and the histone mark may reflect a shared upstream transcriptional program; or *DNAAF3-AS1* expression may simply mark a cell type or proliferative state whose transcriptome, and consequently its H3K36me3 landscape, differs, without any regulatory relationship between the two.

**Figure 4.**
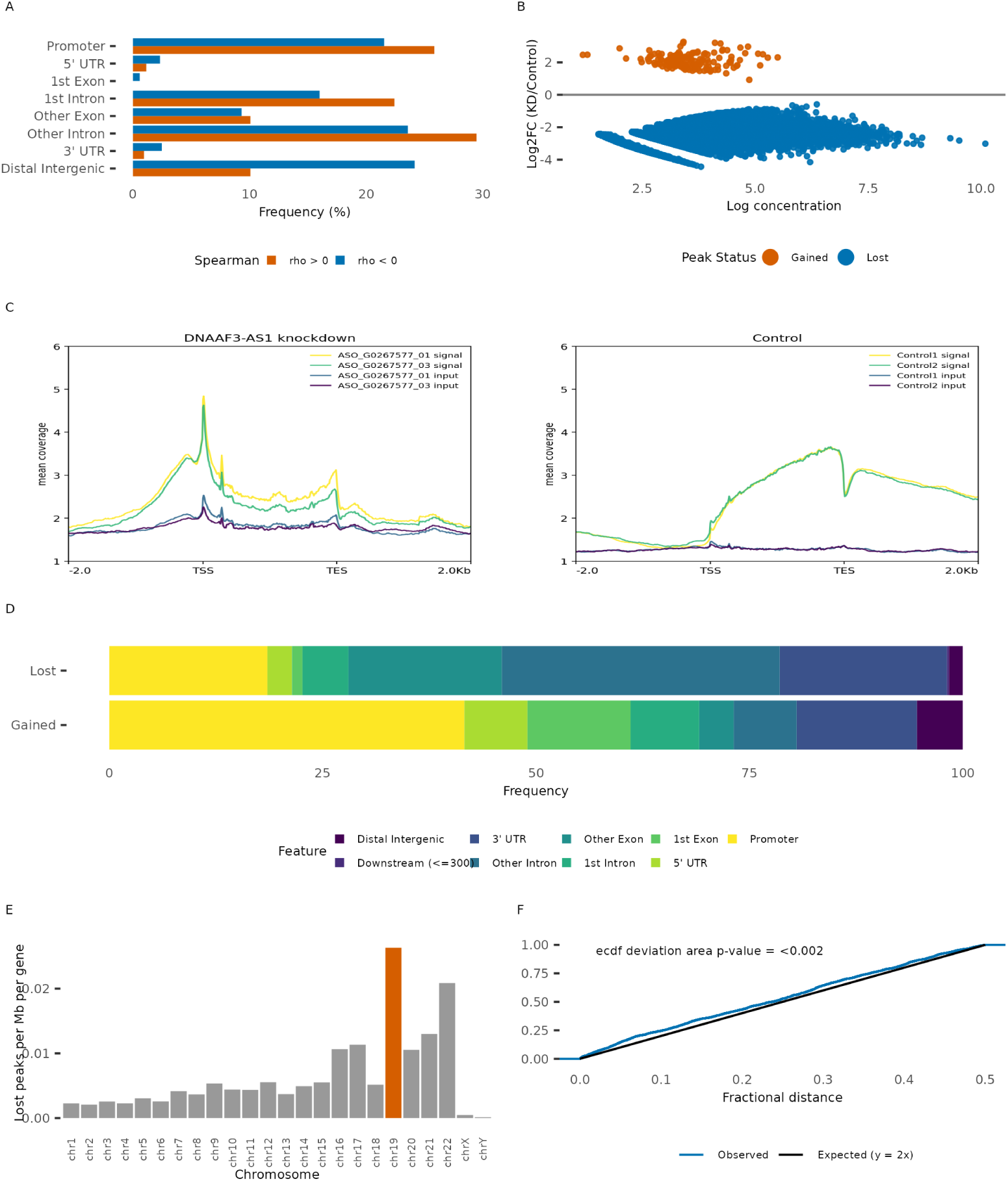
DNAAF3-AS1 knockdown alters the H3K36me3 landscape. **A** Genomic feature distribution of HiMoRNA H3K36me3 peaks associated with DNAAF3-AS1 expression, split by the sign of the Spearman correlation (rho *>*0 vs rho *<*0). **B** Differential H3K36me3 peaks (FDR *<* 0.05): log2 fold change (knockdown/control) versus log concentration, coloured by peak status (Gained, Lost, Unchanged). **C** Metagene profiles of RPKM-normalised H3K36me3 ChIP-seq signal over the set of highly expressed genes (mean CAGE expression across the controls is 50% of the maximum gene expression). **D** Genomic feature annotation of Gained versus Lost H3K36me3 peaks. **E** Lost H3K36me3 peaks (log2FC *<* 0) per chromosome, normalised for chromosome length and gene number. For each chromosome the plotted value is *n*_lost_*/*(*L/*10^6^)*/G*, where *n*_lost_ is the number of lost peaks on that chromosome, *L* its length in base pairs (hg38) and *G* the number of genes on it; chr19, the DNAAF3-AS1 host chromosome, is highlighted in orange. **F** GenometriCorr relative-distance analysis of lost H3K36me3 peaks: empirical cumulative distribution of the observed peak-to-reference relative distances versus the uniform expectation (*y* = 2*x*).

We examined how *DNAAF3-AS1* knockdown affected the expression of H3K36 methyltransferases and demethylases (Figure S4C). For *SETD2*, the main writer of H3K36me3, the expression changes were not concordant between the two ASOs. Changes that were significant and concordant for both ASOs were all decreases: four writers of H3K36me1/me2, *NSD1*, *WHSC1(NSD2)*, *WHSC1L1(NSD3)*, and *ASH1L* [56]; *KDM2B*, a demethylase of H3K36 [25]; and *SETD3*. *SETD3* was originally described as an H3K4/H3K36 methyltransferase [19] but is now established primarily as an actin histidine methyltransferase [31], so the relevance of its downregulation to H3K36 methylation is uncertain. The remaining changes were significant for a single ASO. *SMYD5*, reported to deposit H3K36me3 at promoters [71], was downregulated with ASO_03 only. With ASO_01 only, *KDM4A*, a demethylase of H3K36me3 and H3K9me3 [7], was strongly upregulated, as were the H3K36me1/me2 demethylase *KDM2A* and, weakly, *SETMAR*, whereas the *KDM4A* paralogs *KDM4B* and *KDM4C* and the methyltransferase *SMYD2* were downregulated. Reduced *SETD2* together with increased *KDM4A* would be expected to lower H3K36me3 levels and could contribute to the loss of H3K36me3 upon knockdown. However, neither change was reproduced with ASO_03, so these effects may reflect the higher knockdown efficiency of ASO_01 or an ASO-specific off-target effect.

To test whether *DNAAF3-AS1* loss influences H3K36me3 distribution, we knocked down *DNAAF3-AS1* in fibroblasts using two independent antisense oligonucleotides, with a non-targeting ASO as a control, and performed H3K36me3 ChIP-seq (Figure S5A-D, Supplementary Table 4A-D). The number of H3K36me3 peaks decreased from 25,000–27,000 in controls to approximately 440 in both knockdowns (FDR *<* 0.05). Differential analysis showed loss of H3K36me3 signal at the majority of peaks (FDR *<* 0.05; Figure 4B, Supplementary Table 4E), peaks gaining signal showed smaller effect sizes. Metagene profiles of highly expressed fibroblast genes (above the median expression, top 50% by TPM in FANTOM6) showed the expected gene-body–enriched distribution in controls (Figure 4C). In knockdown samples, this profile was flat, approaching background with spikes in TSS and TES, H3K36me3 retained in the gene bodies of a small number of genes (Figure S6A,B). The largest fraction of the lost peaks was in introns, exons and 3’ UTRs, mirroring the H3K36me3 peak distribution in controls (Figure 4D). Peaks that gained signal upon knockdown were more frequent in promoters compared to the lost peak distribution (41.6% vs 18.5%). The number of peaks lost upon knockdown normalized by chromosome length and the number of genes per chromosome was the greatest on chromosome 19, which is the chromosome of origin for *DNAAF3-AS1* (Figure 4E). This may suggest that the *DNAAF3-AS1* transcript acts in cis.

### DNAAF3-AS1 knockdown ChIP-seq supports apoptotic regulation

To trace the functional consequences of *DNAAF3-AS1* knockdown based on the ChIP-seq results, we analysed pathways that were activated upon knockdown using genes that acquired gene-body H3K36me3 as a mark of activated expression. These targets converged on unfolded protein response, DNA damage tolerance and cGMP–PKG signaling, cell cycle (p-value *<* 0.01, Figure S6C), mirroring the p53 activation signature from FANTOM6. Two competing hypotheses can account for the relationship between this stress response and the genome-wide collapse of H3K36me3. Under the first, the loss of H3K36me3 is a consequence of the stress response: *DNAAF3-AS1* knockdown induces DNA damage and p53 activation through an H3K36me3-independent route, and H3K36me3 is lost secondarily, through RNA polymerase II degradation triggered by DNA repair [13] or through chromatin condensation during apoptosis [45]. Under the second, the loss of H3K36me3 is the primary event: because H3K36me3 engages the DNA repair machinery [58], its depletion would lead to the accumulation of DNA damage and thereby to activation of the p53 response. Our data do not discriminate between these hypotheses. They could be distinguished by combining *DNAAF3-AS1* knockdown with TP53 knockdown or pharmacological p53 inhibition: persistence of the H3K36me3 loss in the absence of p53 activity would place it upstream of the p53 response, whereas its rescue would indicate that it is secondary to p53-dependent apoptosis. A time course resolving the order of H3K36me3 loss, accumulation of DNA damage markers, and p53 stabilisation would additionally test whether H3K36me3 is lost through damage-induced, p53-independent degradation of RNA polymerase II.

### RNA-chromatin contacts support DNAAF3-AS1 role in H3K36me3 regulation

We assessed colocalization between two sets of genomic intervals: H3K36me3 ChIP-seq peaks lost upon knockdown and RNA-chromatin contacts (Figure S7A,B, Supplementary Table 5). Peaks lost upon knockdown colocalized with *DNAAF3-AS1* RNA-chromatin contacts more than expected under spatial independence (permutation p-value *<* 0.02, Figure S7C). This association remained significant when chr19 was excluded (permutation p-value *<* 0.002, Figure 4F, Figure S7D), indicating that it is not driven solely by *DNAAF3-AS1* contacts with its own locus during transcription. The non-random association indicates that *DNAAF3-AS1* contacts and knockdown-sensitive H3K36me3 regions occupy overlapping genomic territory, compatible with a direct role for *DNAAF3-AS1* at these sites.

### DNAAF3-AS1 potential role in spermatogenesis

As *DNAAF3-AS1* showed its highest expression in testis, we examined its expression across the finer-grained cell types of this tissue (Figure 5A, Figure S8A-C). Cells from the adult human testis single-cell atlas [22] were annotated by cell type, with sperm cells forming three clusters along a continuum at the distal end of the germline trajectory: Sperm1 retained residual expression of the acrosomal gene ZPBP, which was lost in Sperm2 and Sperm3, whereas Sperm3 showed the highest PRM2 and reduced TNP1 levels. *DNAAF3* expression increases at the transition from early to late primary spermatocytes and is maintained through the subsequent stages (Figure 5B). *DNAAF3-AS1* expression begins to rise in round spermatids and reaches its highest levels in elongated spermatids and the Sperm1 cluster, where it was detected in 37.1% and 32.1% of cells, respectively (Supplementary Table 6).

**Figure 5.**
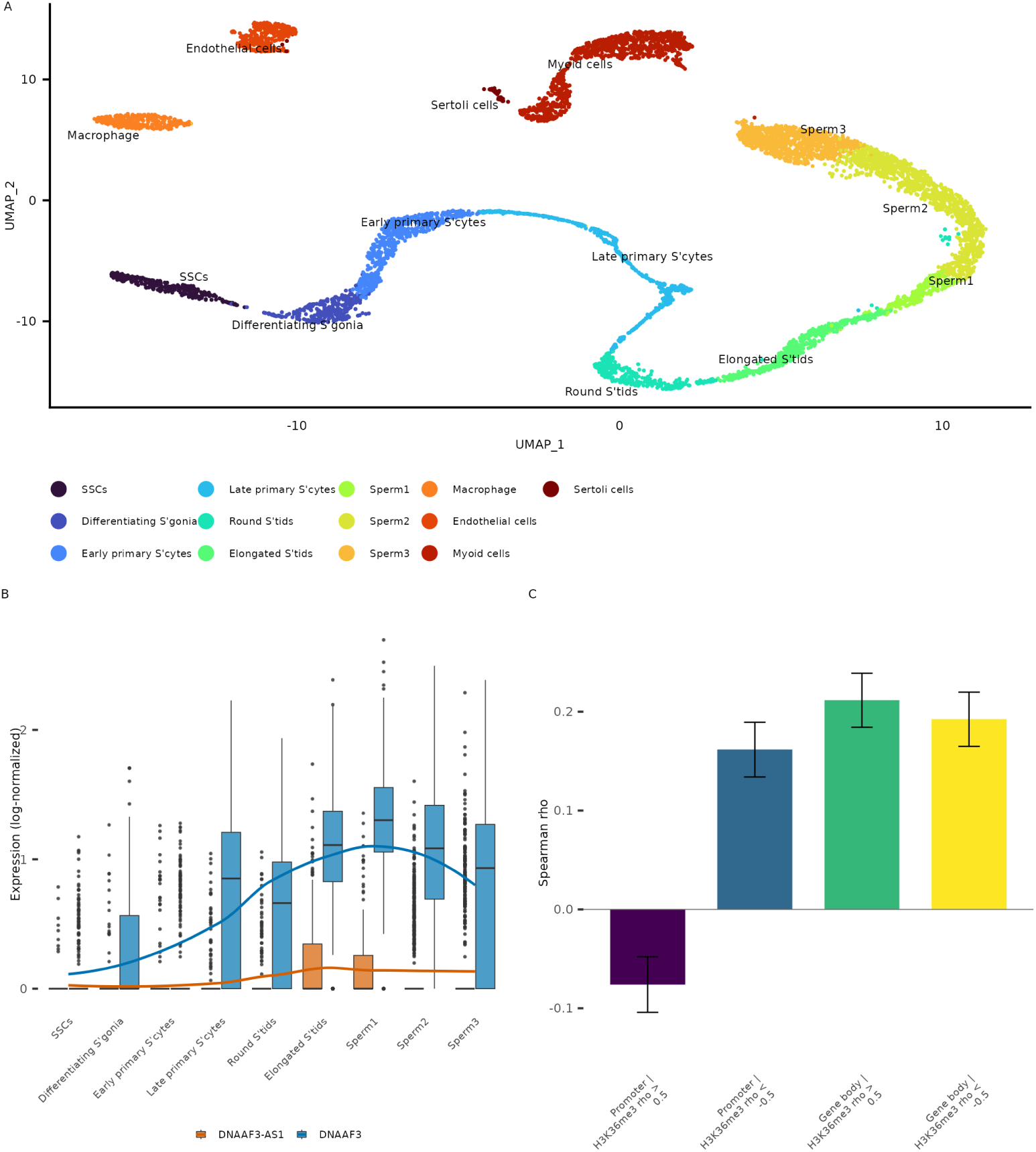
DNAAF3-AS1 expression and HiMoRNA module activity across spermatogenesis stages. **A** UMAP of the single-cell testis dataset (GSE112013), coloured by annotated cell type. **B** Log-normalised expression of DNAAF3-AS1 and DNAAF3 across germ cell types. **C** Spearman correlation between DNAAF3-AS1 expression and the H3K36me3 HiMoRNA module score across all germ cells (n=4817), for each of the four gene sets defined by genomic compartment (promoter versus gene body) and direction of HiMoRNA correlation (rho *>* 0.5 versus rho *<* 0.5).

Histones are largely replaced by protamines during the late stages of spermatogenesis [63]; however,H3K36me3 is retained throughout all spermatocyte stages and in round spermatids, with elongated spermatids showing weaker signal [76]. To test whether the influence of *DNAAF3-AS1* on H3K36me3 distribution could affect gene expression in these cell types, we examined how genes whose H3K36me3 ChIP-seq signal correlates with *DNAAF3-AS1* expression in the HiMoRNA database behave across spermatogenic stages. We considered two gene sets separately, depending on where the correlated H3K36me3 peaks are located: gene-body peaks and promoter peaks, and, within each, distinguished peaks positively (*>*0.5) and negatively (*<*0.5) correlated with *DNAAF3-AS1* expression in the HiMoRNA database, the analysis was performed in germ cells only. Module scores of both gene-body sets correlated weakly and positively with *DNAAF3-AS1* expression (rho = 0.212, 95% CI [0.184–0.239] for the positively correlated set; rho = 0.192, 95% CI [0.164–0.219] for the negatively correlated set; Figure 5C). Because this association was independent of the sign of the H3K36me3 correlation, it more likely reflects a global transcriptional state, such as the late spermatid programme. In contrast, the promoter sets behaved in a sign-dependent manner: the module score of genes with positively correlated promoter peaks was negatively correlated with *DNAAF3-AS1* expression (rho = -0.076, 95% CI [-0.104—0.048]), whereas that of genes with negatively correlated promoter peaks was positively correlated (rho = 0.162, 95% CI [ 0.134–0.189]) and reached its maximum in elongated spermatids (Figure S8D). This pattern is compatible with an association between *DNAAF3-AS1* expression and promoter-level H3K36me3 during spermatogenesis. However, given the weak effect sizes and the sign-independent behaviour of the gene-body sets, it may equally reflect stage-specific transcriptional programmes. The increase in *DNAAF3-AS1* expression during late spermatogenesis could potentially contribute to the removal of H3K36me3 from promoters and to their activation, but H3K36me3 was not profiled in these cells, and direct experimental testing would be required to establish causality.

## Discussion

*DNAAF3-AS1* is an lncRNA whose expression correlates genome-wide with the intensity of H3K36me3 in the HiMoRNA database. Here, we characterise its functional and epigenetic repertoire. Antisense oligonucleotide knockdown in human dermal fibroblasts, followed by H3K36me3 ChIP-seq, showed that loss of *DNAAF3-AS1* reshapes the H3K36me3 landscape, collapsing gene-body enrichment while redistributing residual signal toward promoters. Signal loss was most pronounced on chromosome 19, the chromosome of origin of *DNAAF3-AS1*, and knockdown-sensitive regions colocalized with *DNAAF3-AS1* RNA-chromatin contacts, together consistent with a direct role in at chromatin rewiring. Independently, transcriptome profiling of *DNAAF3-AS1* knockdown in the FANTOM6 dataset revealed activation of the p53 pathway. Because the transcript reaches its highest expression in testis, we then used the adult human testis atlas [22] to ask whether the same axis might operate during spermatogenesis.

Whether the activation of p53 and the remodelling of H3K36me3 are mechanistically linked is a central question raised by these results. H3K36me3 recruits components of the DNA damage response and of the mismatch and recombination repair machinery to transcribed chromatin, and its depletion impairs damage recognition [38, 58]; loss of the mark could therefore itself contribute to p53 activation. A second open question is why H3K36me3 collapses so extensively after *DNAAF3-AS1* knockdown. Expression of *SETD2* did not change concordantly between the two ASOs, arguing against a simple loss of the enzyme that deposits the mark. Instead, both ASOs downregulated *NSD1*, *NSD2*, *NSD3*, and *ASH1L*, the writers that generate H3K36me1/me2. Conditional loss of Nsd2 in mouse germ cells reduces both H3K36me2 and H3K36me3 in vivo [35], showing that loss of an upstream H3K36me2 writer can lower H3K36me3 levels. However, this evidence is indirect, as H3K36me2 was not measured here, and the downregulation of these writers may itself be secondary to cell-cycle arrest, since *NSD2* expression in particular is linked to proliferation. A more direct candidate is *KDM4A*, a demethylase of H3K36me3, which was induced by ASO_01, although not by ASO_03. The kinetics of the loss also constrain the mechanism. In arrested cells, H3K36me3 is not diluted by replication and given the reported half-life of H3K36me3 of approximately 57 h [74], reduced writer activity alone is unlikely to account for a near-complete loss of the mark within the time frame of the knockdown; such a loss would require active demethylation, for which *KDM4A* is a candidate, or a loss of transcription accompanied by histone turnover. Consistent with the latter, we cannot yet exclude that the genome-wide loss of H3K36me3 is secondary to the stress response itself, through RNA polymerase II degradation during DNA repair or through the global chromatin condensation that accompanies programmed cell death.

One setting in which these observations might be physiologically relevant is the male germ line. *DNAAF3-AS1* is broadly co-expressed with its antisense partner *DNAAF3* across spermatogenic stages, and the promoter-level H3K36me3 programme we identified tracks spermatogenesis stages. In mouse germ cells, loss of Nsd2 reduces H3K36me2 and H3K36me3 and causes both spermatocyte apoptosis and aberrant spermiogenesis, with impaired histone removal [35]. Because *NSD2* was downregulated with both ASOs in the FANTOM6 data, one possibility is that *DNAAF3-AS1* contributes to the regulation of *NSD2* during spermatogenesis. As *DNAAF3-AS1* expression rises only after meiosis, such a role would be confined to round and early elongating spermatids, where H3K36me3 is still present, and would relate to the spermiogenic rather than the meiotic function of *NSD2*. This hypothesis extrapolates from fibroblasts, in which reduced *NSD2* expression may be secondary to cell-cycle arrest, and remains to be tested in germ cells.

Several limitations temper these interpretations. The mechanism was studied in dermal fibroblasts, a cell type distant from the testis. The germ-cell analysis is correlational and derived from a single-cell atlas rather than from perturbation. Testing this model directly will require perturbing *DNAAF3-AS1* in a germ-cell and time-resolved, spike-in-normalised H3K36me2/me3 ChIP-seq to establish whether substrate depletion of H3K36me2 accompanies H3K36me3 loss, and rescue of *NSD2* or of H3K36me2 to test whether it restores H3K36me3 and averts apoptosis. Mapping the direct RNA-protein and RNA-chromatin partners of *DNAAF3-AS1* would further clarify whether it scaffolds the H3K36 methylation machinery.

## Data Availability

The ChIP-seq next-generation sequencing data (unprocessed read files) can be accessed under project accession PRJNA1551676. The scripts used to analyze data can be accessed at https://github.com/MedvedevaLab/DNAAF3-AS1.

## Supplementary Data statement

## Acknowledgements

Author Contributions Statement

## Supplementary Information

**Figure S1.**
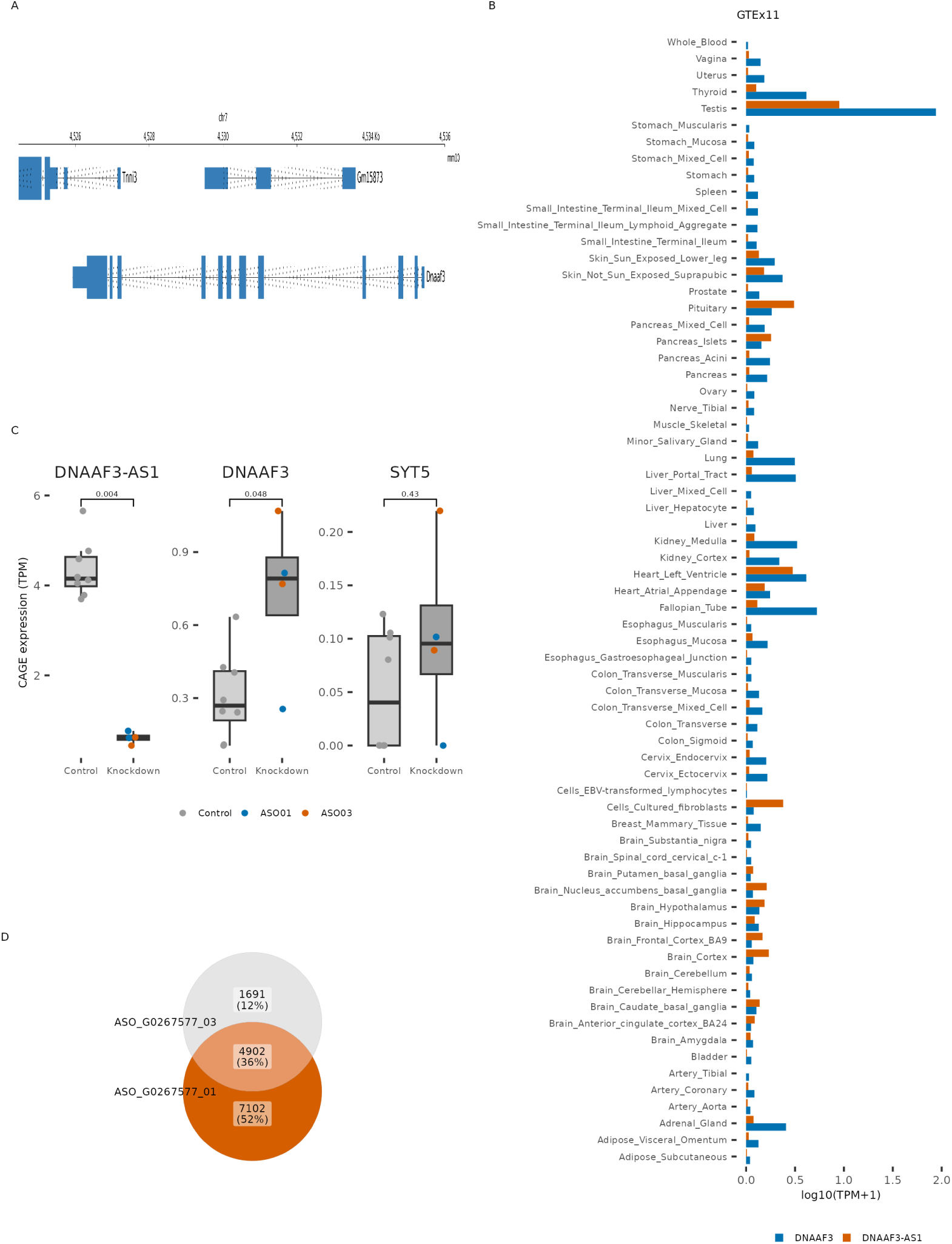
**A** Genomic tracks of the Gm15873 in mouse genome mm10. **B** Median expression (log10(TPM+1)) of DNAAF3-AS1 and DNAAF3 across GTEx v11 tissues. **C** FANTOM6 CAGE expression (TPM) of DNAAF3-AS1, DNAAF3 and SYT5 in control versus DNAAF3-AS1-knockdown, p-values are from two-sided Wilcoxon rank-sum tests (control versus knockdown). **D** Overlap of significantly differentially expressed genes (FDR *<* 0.05) between ASO_01 and ASO_03.

**Figure S2.**
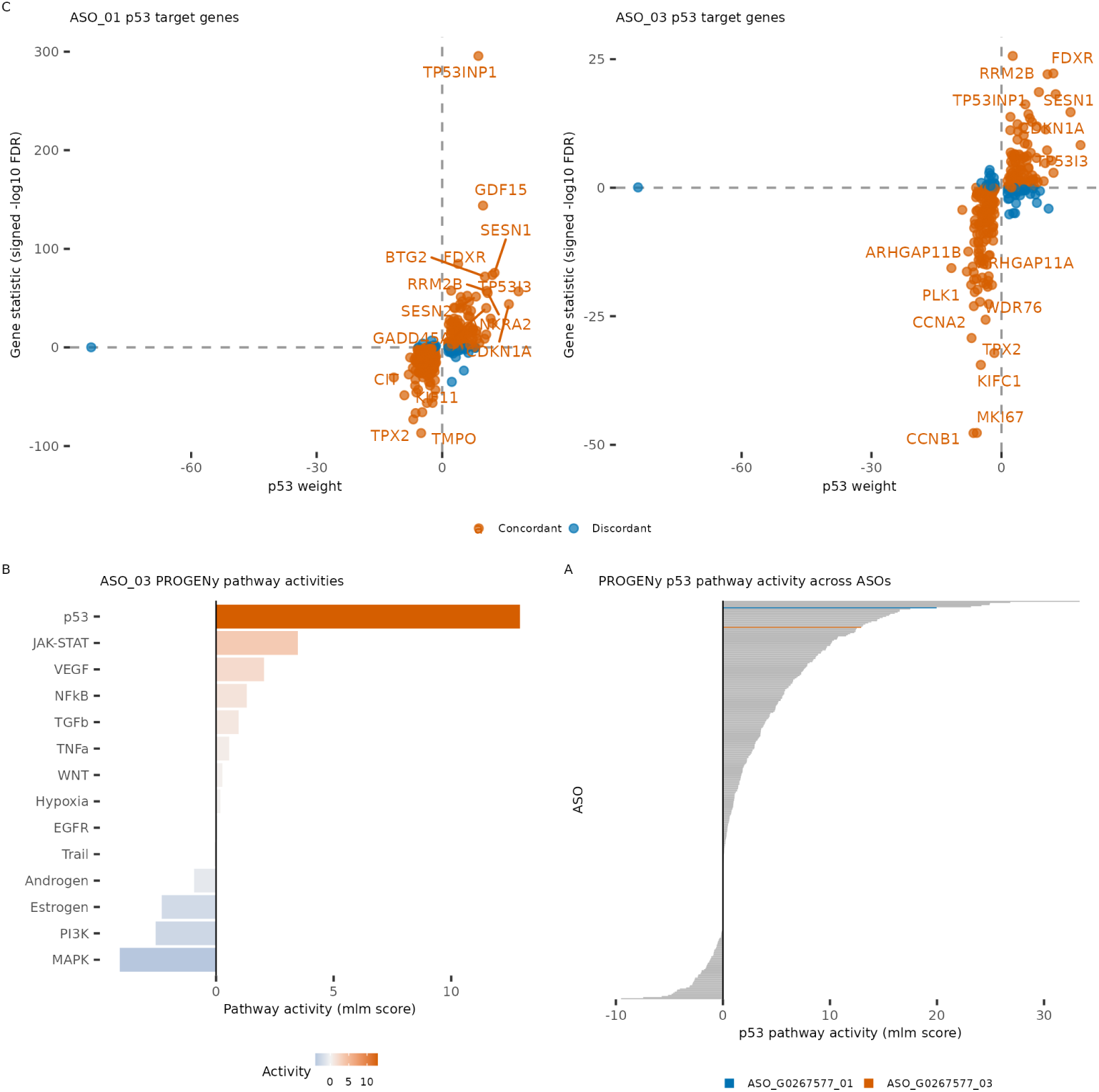
**A** PROGENy pathway activities inferred with decoupleR for the ASO_03 differential expression signature. **B** decoupleR p53 target gene activities for ASO_01 (left) and ASO_03 (right). **C** PROGENy p53 pathway activity across all ASOs in FANTOM6.

**Figure S3.**
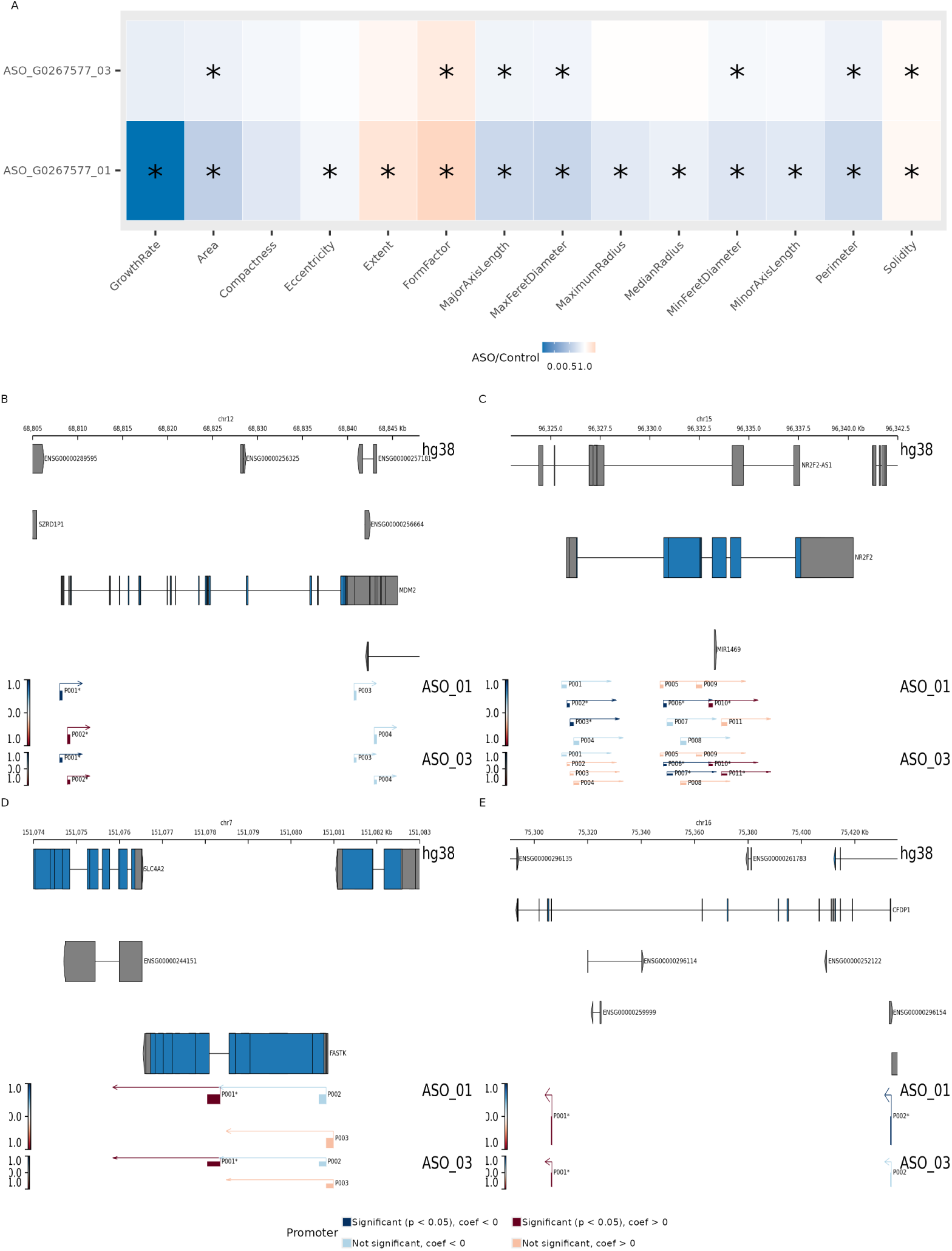
**A** CellProfiler cell morphology features normalized by NCA ASO transfection for ASO_01 and ASO_03 **B-E** Genomic tracks at the MDM2 (B), NR2F2 (C), FASTK (D) and CFDP1 (E) loci showing differential promoter usage upon knockdown with ASO_01 and ASO_03; asterisks mark promoters with significant differential usage (prmtrPv *<* 0.05).

**Figure S4.**
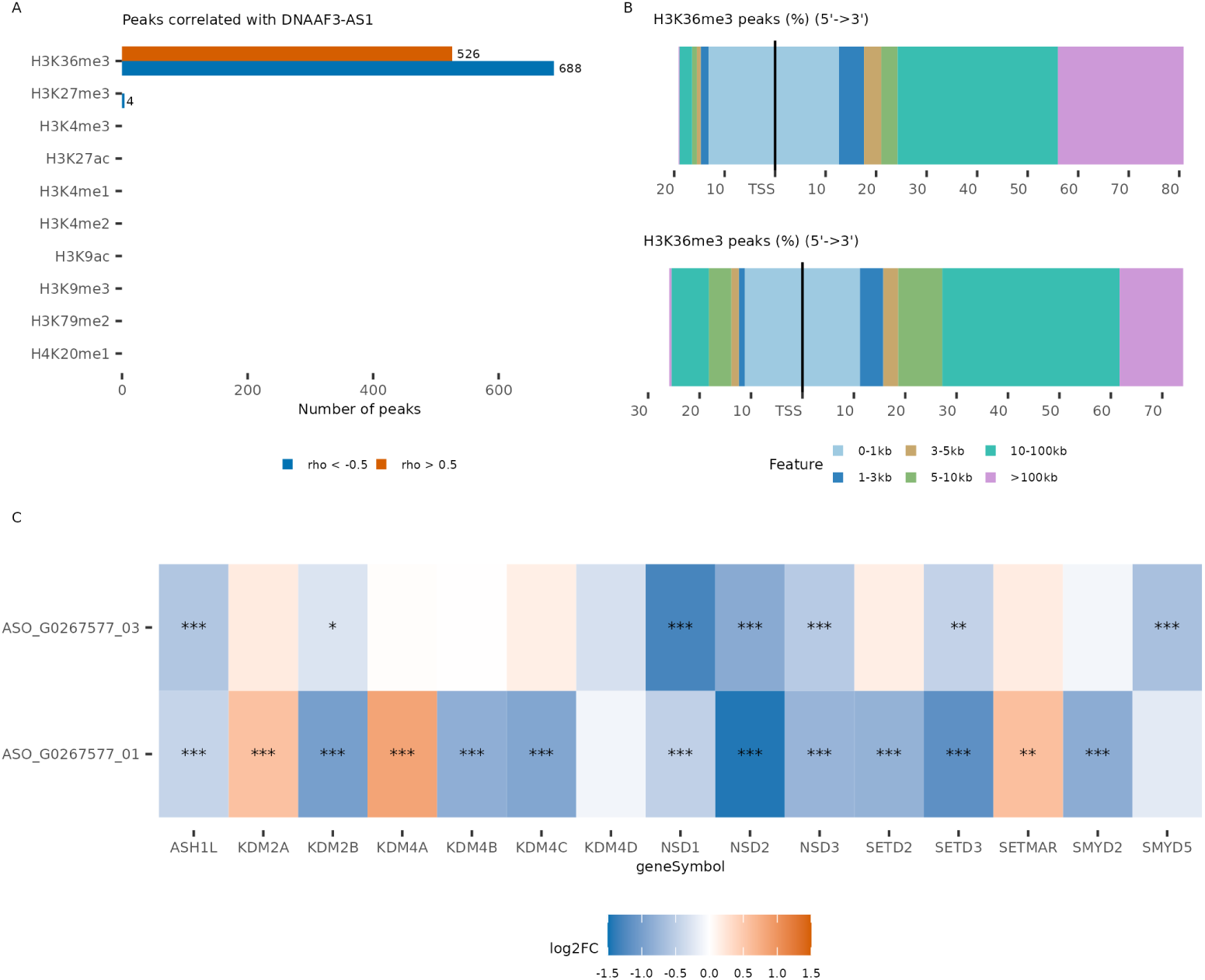
**A** Number of HiMoRNA peaks correlated with DNAAF3-AS1 expression per histone mark. **B** Distribution of H3K36me3 HiMoRNA peaks relative to the transcription start site. **C** Differential expression (log2FC) of H3K36 methyltransferase and demethylase genes upon DNAAF3-AS1 knockdown, asterisks denote significance (* FDR *<* 0.05, ** FDR *<* 0.01, *** FDR *<* 0.001).

**Figure S5.**
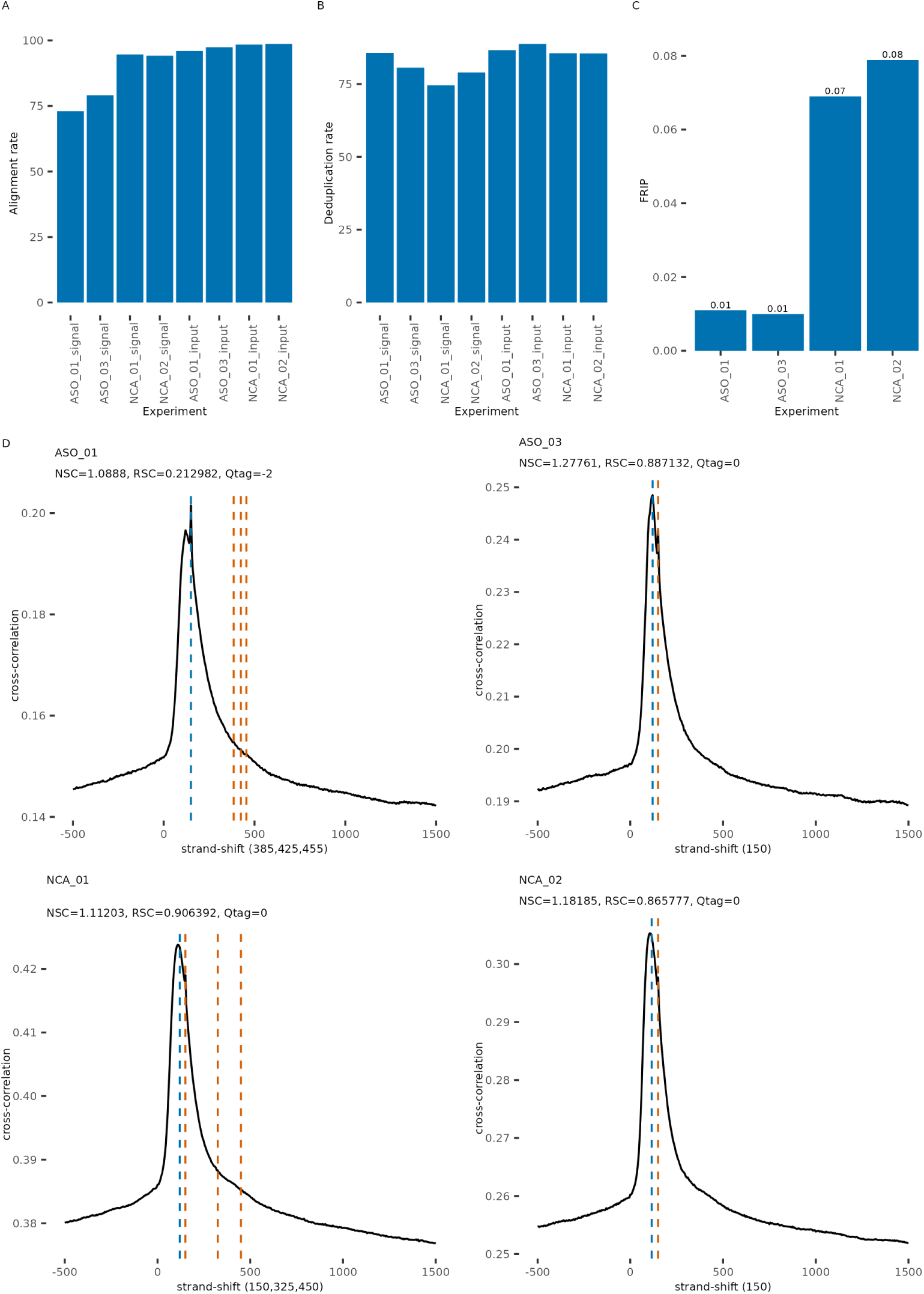
**A-C** Alignment rate (A), deduplication rate (B) and fraction of reads in peaks (C) for each ChIP-seq library. **D** Strand cross-correlation profiles for ASO_01, ASO_03 and controls; dashed lines mark the ChIP cross-correlation peak (red) and the phantom (read-length) peak (blue).

**Figure S6.**
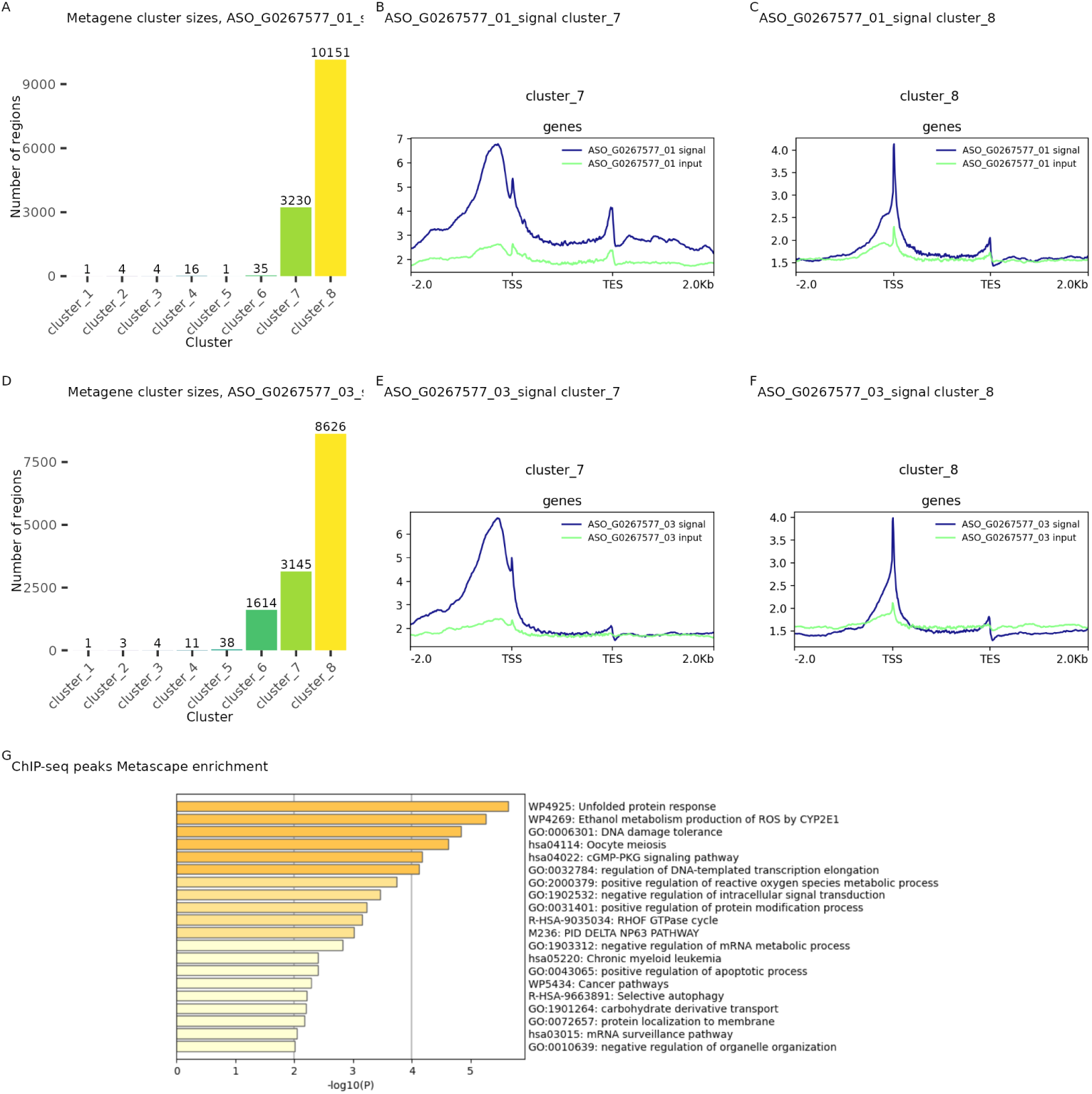
**A** Sizes of the metagene clusters for the ASO_01 signal sample. **B, C** Metagene profiles of clusters 7 (B) and 8 (C) for the ASO_01 signal sample. **D** Sizes of the metagene clusters for the ASO_03 signal sample. **E, F** Metagene profiles of clusters 7 (E) and 8 (F) for the ASO_03 signal sample. **G** Metascape functional enrichment analysis of the genes associated with the ChIP-seq peaks gained upon knockdown.

**Figure S7.**
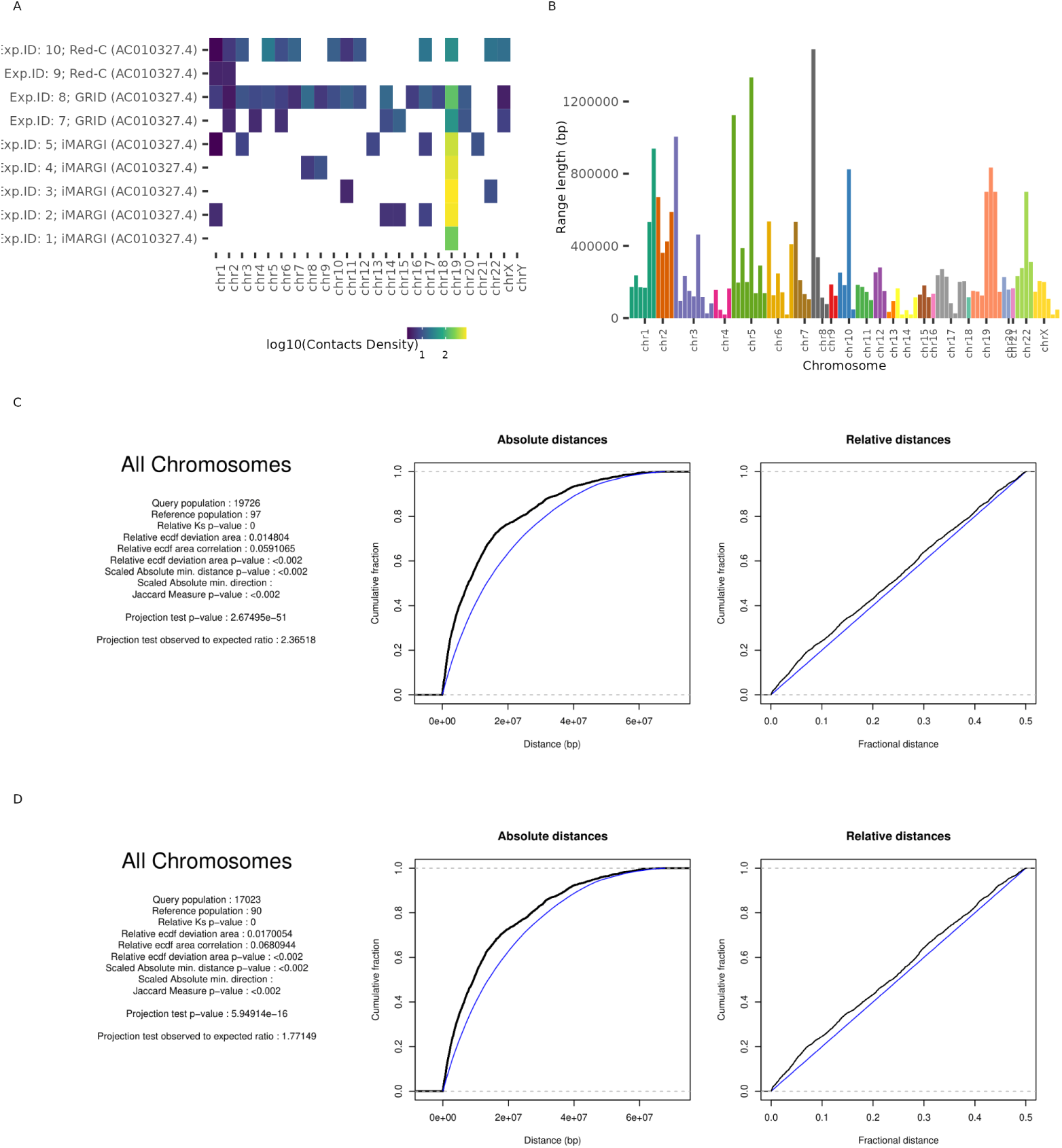
**A** Density of DNAAF3-AS1 RNA–chromatin contacts across chromosomes (log10 scale) for each RNA–chromatin interaction experiment in the RNAChromDB dataset (iMARGI, GRID and Red-C); missing values are shown in white. **B** Length (bp) of each DNAAF3-AS1 contact cluster. **C** GenometriCorr spatial-correlation report between Lost H3K36me3 peaks (DiffBind Fold *<* 0) and DNAAF3-AS1 RNA–chromatin contact clusters (all chromosomes). **D** As in C, after excluding the DNAAF3-AS1 host chromosome (chr19).

**Figure S8.**
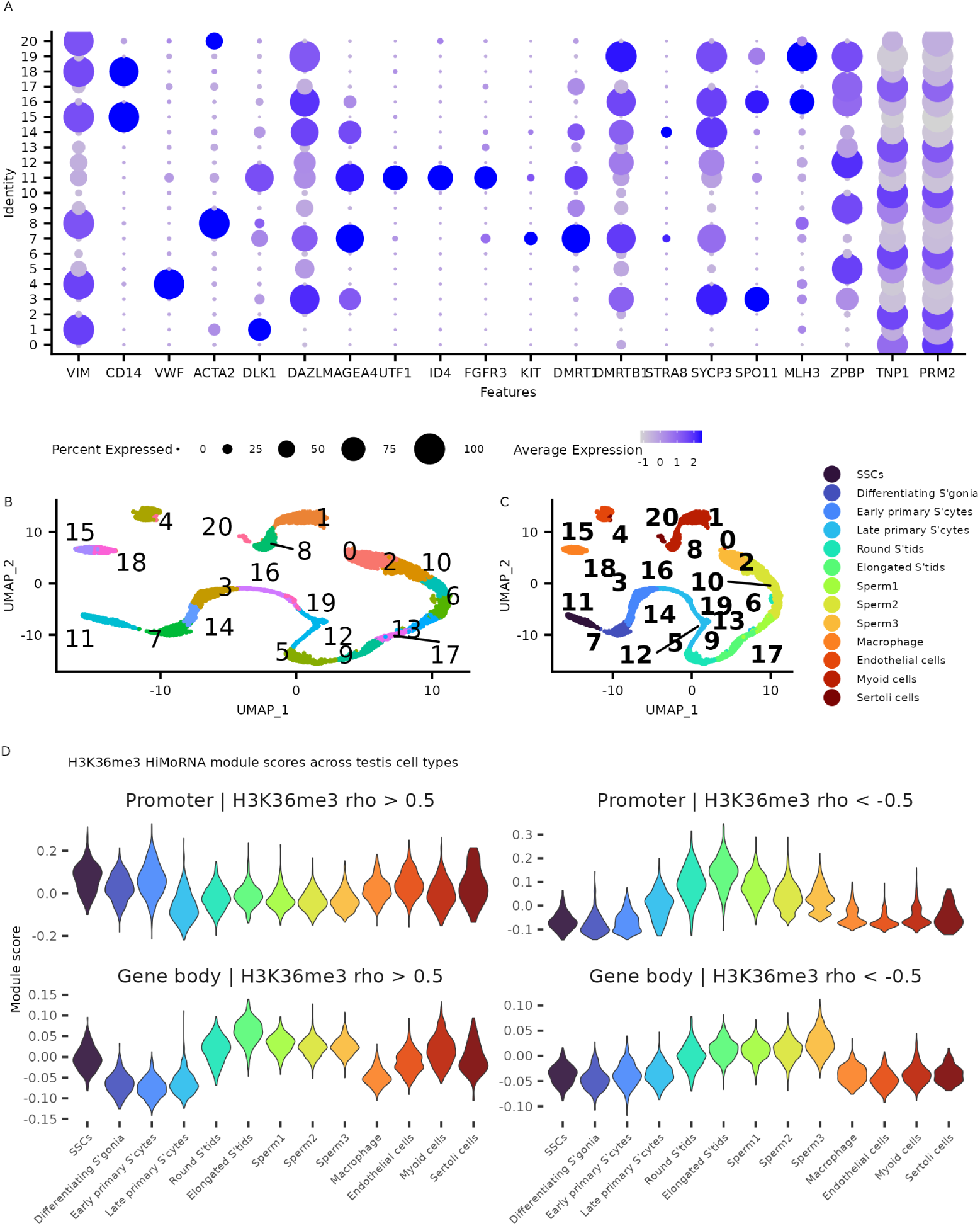
**A** Dot plot of canonical marker genes across the Seurat clusters of the single-cell testis dataset (GSE112013). **B,C** UMAP plots coloured and labelled by Seurat cluster (B) and annotated cell type (C). **D** Violin plots of H3K36me3 HiMoRNA module scores across testis cell types, for the four gene sets defined by genomic compartment and correlation sign.

